# snpXplorer: haplotype-block representation of GWAS signals for integrated variant and cross-trait interpretation

**DOI:** 10.64898/2026.07.20.739485

**Authors:** Niccolò Tesi, Gilad Green, Alex Salazar, Sven van der Lee, Marc Hulsman, Henne Holstege, Marcel Reinders

## Abstract

Genome-wide association studies (GWAS) have identified thousands of loci associated with complex traits, yet translating these signals into biological insight remains challenging: most associated variants are non-coding and lie within linkage disequilibrium (LD) blocks, so interpretation requires functional and cross-trait evidence assembled within haplotypic context. We present an extended version of snpXplorer, an interactive web platform that represents GWAS signals at the haplotype-block level rather than at the variant level. Haplotype-blocks are precomputed genome-wide by partitioning the genome on recombination rate and clustering variants by pairwise LD across European or African ancestries, aggregating association signals, annotations and phenotype associations onto inherited units of variation. The platform incorporates more than 11,000 European and African GWAS datasets from OpenGWAS and enables multi-scale analysis across variants, haplotype-blocks, genes and traits, integrating clinical annotations (ClinVar), allele frequencies (gnomAD), functional predictions (CADD, AlphaGenome), quantitative trait loci (GTEx), structural variation and reported GWAS associations within a single unified interface. Redundancy across closely related phenotypes is reduced by clustering trait descriptions with pretrained biomedical language embeddings, making navigation across large GWAS repositories tractable. Applied to Alzheimer’s disease, snpXplorer identified a haplotype-block at the *TMEM106B* locus linked to eight distinct traits, revealing synergistic pleiotropy across neurological and behavioral phenotypes alongside antagonistic pleiotropy with height. The platform additionally exports PRS-ready variant sets, at user-selected p-value thresholds, for every ingested GWAS. By combining haplotype-aware representation, integrated variant annotation and cross-trait exploration, snpXplorer reduces the need for fragmented queries across databases and lowers the barrier to biological interpretation of GWAS results.

## Introduction

Genome-wide association studies (GWAS) have transformed our understanding of the genetic architecture of complex traits and diseases by identifying thousands of associated loci. However, identifying the genes and biological mechanisms underlying GWAS loci remains a major challenge. Most GWAS signals map to non-coding regions of the genome and encompass multiple correlated variants in linkage disequilibrium (LD), making it difficult to identify the causal variant or gene. These groups of linked variants, representing haplotype-blocks, may provide more biologically meaningful units than individual variants by capturing the combined effects of inherited genetic variation, and may help prioritize the genes most likely affected by the associated haplotype. In addition to single nucleotide variants, disease-associated haplotypes may also harbor larger structural variants, such as transposable elements or tandem repeats, which can further contribute to regulatory and disease mechanisms [1,2].

At the same time, the growing availability of GWAS summary statistics across thousands of traits has also enabled systematic cross-trait analyses. However, these resources contain substantial redundancy, as closely related phenotypes often share overlapping association signals. Efficient exploration therefore requires tools capable of integrating evidence across studies while reducing complexity and preserving biological context.

In 2021, we introduced snpXplorer [3], a web server designed to support regional visualization and functional annotation of SNP data. Since its initial release, snpXplorer has been widely adopted by the genetics community. Motivated by user feedback, advances in large-scale GWAS resources and variant annotation databases, as well as the increasing relevance of haplotype-based analyses, we have developed a substantially expanded version of the platform. Compared with the original 2021 release, the updated snpXplorer extends beyond regional SNP visualization and batch annotation to support large-scale cross-trait GWAS exploration, haplotype-block-level aggregation of association signals, integrated single-variant interpretation, semantic phenotype clustering, and PRS-oriented workflows. These additions reposition snpXplorer from a regional annotation server toward a broader framework for interactive interpretation of complex disease genetics.

Here, we describe how snpXplorer places GWAS association signals in haplotype context, and apply it to Alzheimer’s disease genetics. Using Alzheimer’s disease (AD) as a test case, we resolve the association signal into haplotype-blocks and show that this representation exposes cross-trait pleiotropy at the *TMEM106B* locus and prioritizes *PLEKHA1* at the chromosome 10q26 locus. We describe the design of the snpXplorer web server that makes these analyses possible across trait-centric, gene-centric, and variant-centric workflows.

## Materials and Methods

### Overview

snpXplorer is an integrated, interactive platform for GWAS exploration and annotation (*Figure 1*), comprising different modules that enable real-time, point-and-click analysis across variants, haplotype-blocks, genes, and traits.

**Figure 1:**
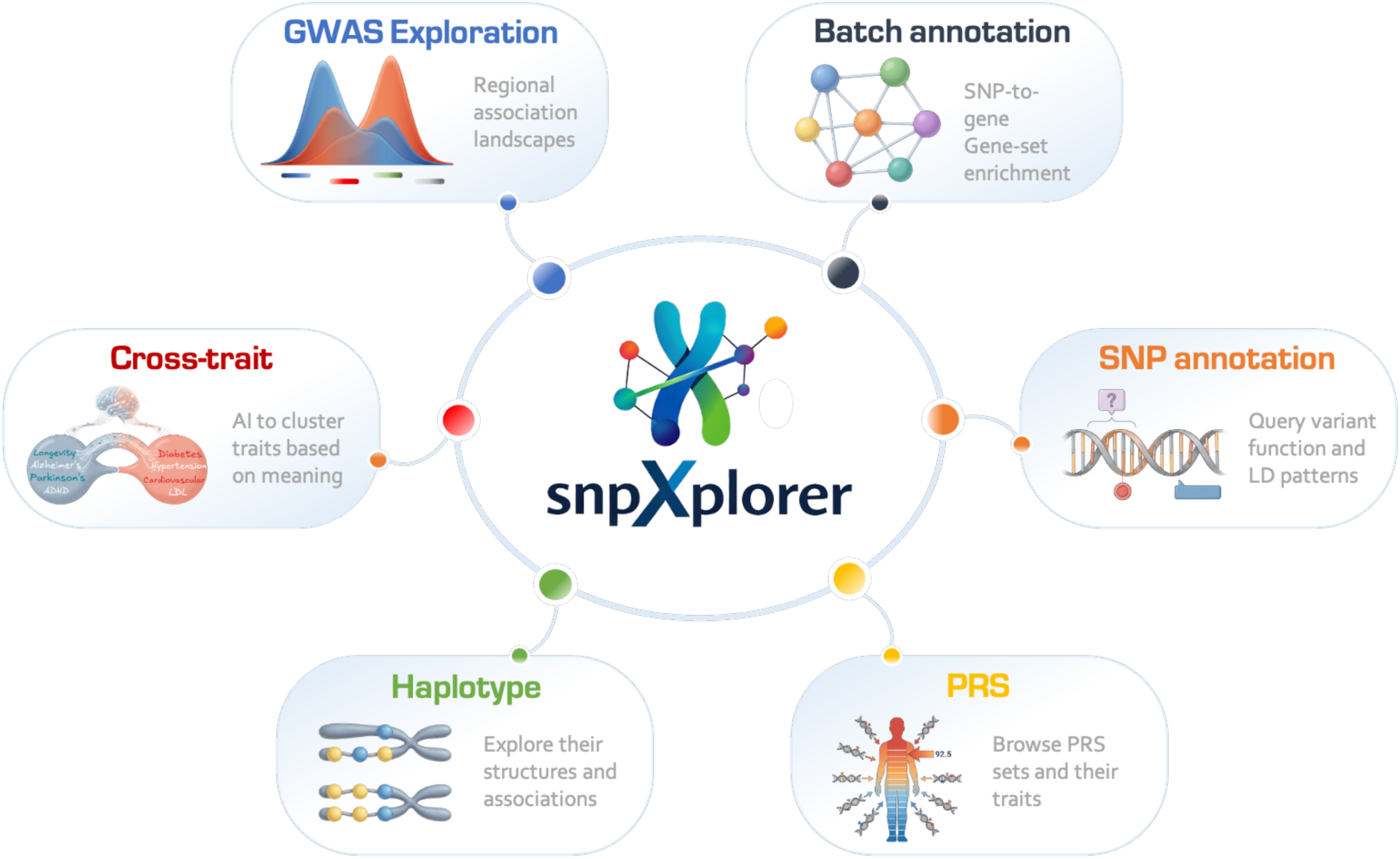
snpXplorer platform. SnpXplorer is a web-based platform freely available at https://snpxplorer.net that allows users to (i) browse *GWAS exploration*: explore, overlay and compare results from >10,000 GWAS studies based on OpenGWAS; (ii) run *Batch annotation*: perform batch annotation of up to 10,000 SNPs to their likely affected genes and optionally gene-set enrichment analysis; (iii) check *SNP annotation*: run single annotation of SNPs to investigate their allele frequencies, clinical evidence, QTLs, predicted functional consequences, LD haplotypes, structural variants, and linked GWAS phenotypes; (iv) explore *Haplotype* structures, the underlying SNPs, and the haplotype effect on human phenotypes from GWAS; (v) identify *Cross-trait* interactions present in OpenGWAS using a semantic similarity-based clustering to simplify redundancy; and (vi) support *PRS* generation providing PRS-ready SNP-sets for all OpenGWAS studies.

### GWAS exploration

The original *GWAS exploration* module supported interactive visualization of a limited set of GWAS association signals across genomic regions of interest. We have now integrated the IEU OpenGWAS database [4,5], enabling users to browse and overlay association signals from over 10,000 GWAS studies from both European and African ancestry without uploading local summary statistics. Users can visualize multiple traits simultaneously, explore LD patterns (defined in a fast-accessing database based on 4,777 European individuals [6], and 820 African individuals [7]), inspect recombination rates and display structural variants [8] (*Figure S1*).

### Batch annotation

The *Batch annotation* module annotates up to 10,000 variants to genes integrating clinical evidence from ClinVar [9], genomic position from RefSeq [10], quantitative trait loci (QTLs) from GTEx [11], and functional predictions from CADD [12] and AlphaGenome [13]. Cross-ancestry allele frequencies from gnomAD [14] as well as phenotype associations from OpenGWAS are also reported. The user can select the ancestry for querying OpenGWAS studies and for retrieving LD partners for the query variants. Optionally, gene-set enrichment analysis can be performed with gProfiler [15]. Gene-set enrichment analysis is followed by clustering of significant terms based on semantic similarity, and clusters are annotated using word clouds to provide functional context (*Figure S2A-B*). To provide haplotype context and improve variant annotation, snpXplorer now incorporates the annotation of variants in LD with the input variants. For example, an intergenic variant in LD with a coding variant can be linked to the affected gene through the shared haplotype-block. Users can additionally submit genomic intervals alongside or instead of individual variants: all variants with minor allele frequency >0.01 within a submitted interval are identified and annotated. As the total number of variants per job is capped to 10,000, when intervals are combined with other intervals and/or individual variants, the annotation budget is allocated proportionally across all inputs, with individual variants prioritized. *Batch annotation* jobs can be submitted along with an email address, and results are sent to the user upon completion (*Figure S2A*).

### SNP annotation

An important new component of snpXplorer is the *SNP annotation* module, which provides a unified interface for querying individual variants. The annotation engine is the same as in the *Batch annotation* module. For a given variant, snpXplorer returns allele information, population frequencies, predicted functional impact, nearby affected genes, QTL associations and previously reported associations with representative traits from OpenGWAS (*Figure S3*). Annotations are also extended to variants in LD (R^2^>0.6, which the user can choose to rely on European or African ancestry) with the query variant, often improving the identification of the candidate affected gene(s). By aggregating heterogeneous sources of evidence into a single view, this module reduces the need for manual cross-referencing across databases and facilitates rapid variant interpretation, supported by interactive tables, graphs and cross-references, and with the possibility to download all content.

### Haplotype-block exploration

The *Haplotypes* module is a new component of snpXplorer. It enables users to explore GWAS signals at the haplotype-block level in both European and African ancestry. Haplotypes are precalculated in a two-step procedure (*Figure S4*). The genome is first split into subregions based on recombination rates (based on HapMap, downloaded from UCSC genome browser, threshold=25) [16,17]. Then, pairwise LD matrices are calculated for all variants in each subregion, using genotyping data from 4,777 unrelated and European individuals (an in-depth description of these individuals is available elsewhere [6]) or 820 African individuals from the 1000Genome project (a substantially smaller and more heterogeneous reference panel than for European ancestry) [7]. Variants are finally clustered into haplotype-blocks by complete-linkage hierarchical clustering, using 1-LD (R^2^ values) as distance measure, with clusters designed to ensure all pairwise LD values are R^2^ values >0.2. This threshold was chosen in line with common convention for defining meaningful linkage disequilibrium, and was further supported by a sensitivity analysis across a range of R^2^ values (*Figure S5).* Haplotype-blocks can be queried by gene, or by individual variant. When querying by gene, all haplotype-blocks starting within 50 kb of the gene’s transcription start site are shown. When querying by variant, all haplotype-blocks starting within 100 kb of the variant position are shown. The platform supports interactive inspection of haplotype-block composition, LD structure, and cross-trait associations, and all resulting tables and graphs can be downloaded.

### Cross-trait exploration

In addition to supporting haplotype-blocks, the *Haplotypes* module allows users to browse phenotypes across large GWAS repositories. All OpenGWAS metadata were obtained, comprising 50,056 studies in total. For European ancestry, we restricted to European studies with a sample size >10,000 (N= 25,217), and further excluded eQTL studies, leading to 10,604 datasets. For African ancestry, we restricted to African studies with a sample size >5,000 (N=751). To reduce redundancy among closely related traits, semantic similarity was approximated using pretrained biomedical language embeddings (BioLORD-2023 [18], SapBERT [19]) applied exclusively to OpenGWAS trait titles (the *trait* field in the downloaded association files), separately for European and African studies. Titles were normalised by lowercasing, replacing selected punctuation with spaces, removing predefined boilerplate phrases (e.g. *GWAS of* and *genome-wide association study of*) and common stop-words, and collapsing repeated whitespace. No additional phenotype descriptions, ontology annotations, or study metadata were included in the embedding input. Embeddings produced by each model were independently L2-normalised and combined using weighted concatenation, with weights of 0.70 for BioLORD-2023 and 0.30 for SapBERT. Pairwise cosine distances between trait embeddings were computed and used as input for agglomerative hierarchical clustering with average linkage. Clusters are defined by applying a cosine distance threshold of 0.5, such that traits within a cluster exhibit high semantic proximity. This threshold was chosen to group only phenotypes with close semantic correspondence rather than maximizing the reduction in redundancy. The number of clusters and traits obtained across cosine distance thresholds ranging from 0.1 to 0.7, together with the UMAP relative to the chosen threshold (0.5), are reported in *Figure S6*. Singleton traits were reassigned to their nearest non-singleton cluster when cosine similarity was at least 0.4. For each cluster, a representative study is selected based on study characteristics such as sample size, number of significant hits, data availability, and data completeness. All studies within a cluster remain accessible: users can view the full list of member studies, including ancestry, sample size, and PubMed identifiers, alongside the selected representative. To identify neighboring clusters, inter-cluster cosine distances are calculated based on the average pairwise distance between traits across clusters. Because both European and African trait descriptions were embedded with the same models, the relative embedding coordinates are comparable. When users query a phenotype of interest, an interactive UMAP embedding showing the cluster containing the phenotype of interest as well as the five closest clusters is displayed, along with the cosine similarity heatmap between all phenotypes. Users can query traits of either European or African ancestry: depending on the ancestry of the queried trait, the user can choose to display both the individual-ancestry UMAP, or the combined European+African UMAP. This strategy is intended to facilitate scalable navigation across large GWAS repositories rather than replace formal genetic correlation analyses.

### Haplotype-block-level phenotype aggregation

For each haplotype-block, phenotype associations are derived by aggregating the effect sizes of its constituent SNPs across the representative traits retained after semantic similarity-based clustering of OpenGWAS phenotypes (see *Cross-trait exploration* above). SNP effect sizes across phenotypes were harmonized by OpenGWAS and further verified by us, ensuring that the effect estimates always referred to the non-reference allele of the human genome. The average effect of haplotype-blocks on representative traits is computed over only SNPs reaching suggestive significance for the trait in question (p<5×10^−5^), using an unweighted mean of per-SNP effect sizes. Because OpenGWAS reports effect sizes on different scales depending on trait type (*e.g.* log odds ratios for binary traits, beta coefficients for continuous traits), effect sizes can be optionally standardized to z-scores prior to averaging. Users can select a specific haplotype-block of interest and inspect its structure, trait associations, and trait-by-trait correlations. The latter is calculated across SNPs constituting the haplotype-block that show suggestive association with each pair of traits being compared, requiring a minimum of two overlapping SNPs per trait pair. Both Pearson and Spearman correlation coefficients can be used, the latter offering robustness to heterogeneity in effect-size distributions across traits. As SNPs within a haplotype-block are, by construction, in strong pairwise LD, these correlations are best interpreted as a summary of how consistently the haplotype-block associate across traits, rather than as an independent-samples statistical test.

### PRS module

Polygenic risk scores (PRS) are weighted scores quantifying an individual’s genetic risk for a given phenotype. The new PRS module provides PRS-ready variant sets for every trait in the OpenGWAS database after QC (see *Cross-trait exploration*). PRS calculation requires a set of variants, the corresponding effect sizes from a GWAS, and individual-level genotyping data; snpXplorer supplies the first two. For any European or African trait in the OpenGWAS database, users can interactively select and download pre-calculated variant sets, at the p-value threshold of interest, to generate a PRS. Individual-level genotyping data are deliberately kept outside the platform, as sharing or uploading such data is restricted by privacy regulations, so PRS calculation itself is performed locally by the user. Variant sets are derived from curated trait definitions or automated clumping-and-thresholding procedures (LD R^2^=0.05, window=500 kb, European or African individuals), followed by conditional analysis [20] (implemented in GWASum, https://github.com/holstegelab/GWASum). The intended scoring model is an additive PRS, computed as the sum of each variant’s dosage of the effect allele weighted by its GWAS effect size. Effect alleles are harmonized from OpenGWAS to the forward strand of the reference genome (GRCh38). Downloaded variant sets include, for each variant, the effect and other allele, effect size and confidence intervals, p-value, and effect allele frequency. These outputs are designed for compatibility with PRS tools such as Jordan (https://github.com/TesiNicco/jordan) and PolyGenius (https://polygenius.holstegelab.eu).

## Results

### Haplotype-block estimation

Across the genome, snpXplorer defines 372,097 common haplotype-blocks comprising 8,225,502 variants in European individuals, with a median size of 94.4 kb (IQR 32.4-249.4 kb) and a median of 11 variants per haplotype-block (IQR 6-23). Haplotype blocks cover 84.5% of the autosomal reference sequence. Of the 10,604 European GWAS datasets retrieved from OpenGWAS (sample size >10,000), semantic clustering reduced the number of distinct phenotypes to 604 trait clusters (*Table S1*). For African ancestry, snpXplorer identified 984,474 common haplotype blocks comprising 15,404,984 variants, with a median size of 48.5 kb (IQR 19.5-123.2 kb), a median of 10 variants per haplotype block (IQR 6-17) and a coverage of 86.1% of the autosomal reference sequence. The 751 African GWAS datasets retrieved from OpenGWAS (sample size >5,000) reduced to 136 after semantic clustering. The larger number of African haplotypes is a likely consequence of the larger number of SNPs available from sequencing data, as well as the larger haplotypic diversity within African samples [7] (*Table S1*).

### Semantic trait clustering aligned related phenotypes across European and African ancestries

To illustrate cross-ancestry trait clustering directly, we queried *triglycerides*, a trait with sufficient African-ancestry GWAS power, within the *Haplotypes* module of snpXplorer. Sixteen African-ancestry GWAS datasets were retrieved (*Figure 2A, Table S2*), with the representative study conducted in Sub-Saharan African samples (N=24,612, ebi-a-GCST90101747, *Figure 2A*). In the combined European+African UMAP embedding, the African triglycerides cluster projected closest to related European lipid clusters, including triglycerides, cholesterol, fatty acids, and phosphoglycerides (*Figure 2B, Table S2*), indicating that semantically related phenotypes cluster together across ancestries despite being embedded independently of each other.

**Figure 2:**
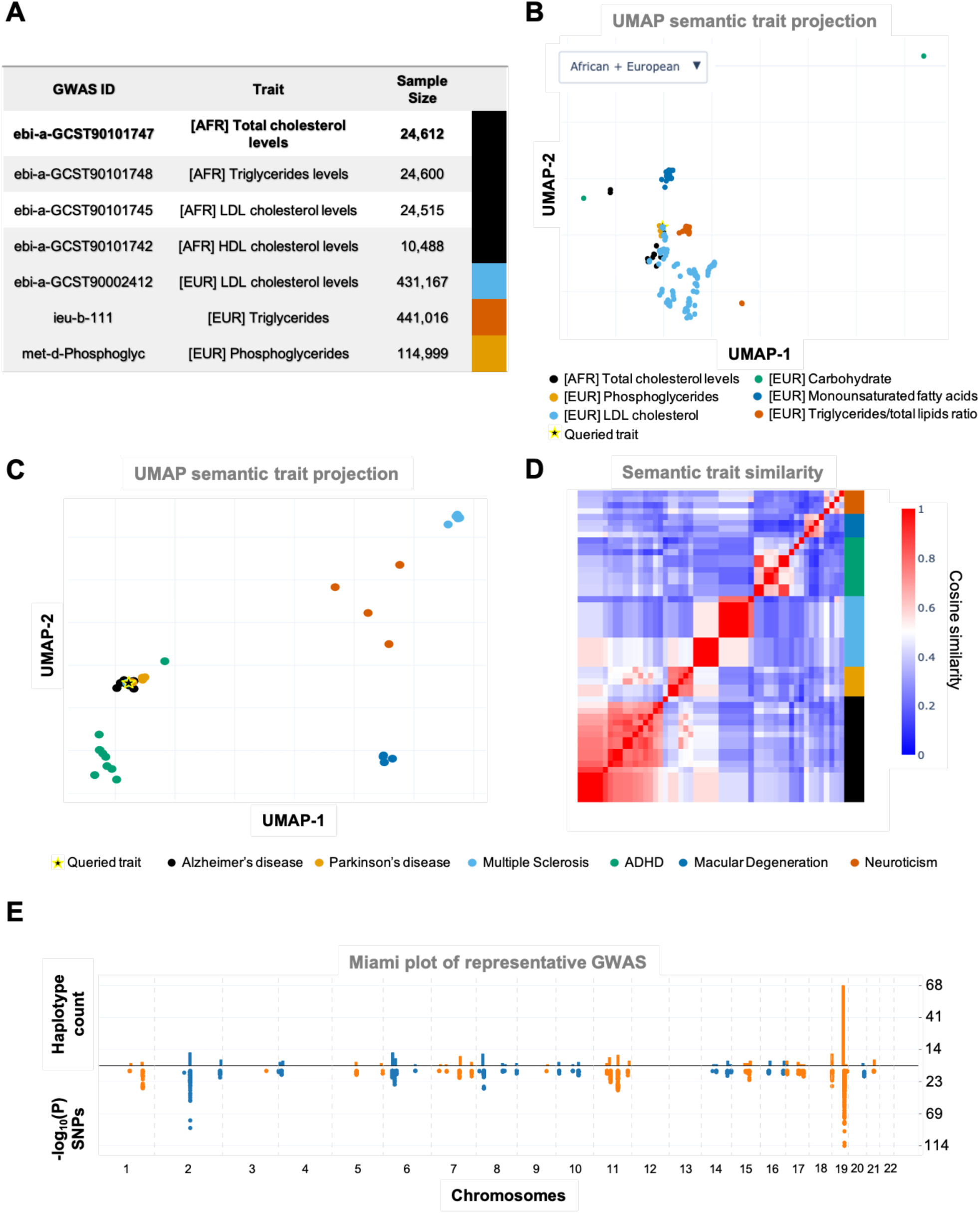
Trait-centric *Haplotypes* module in snpXplorer. (A) Summary table reporting the GWAS from OpenGWAS of African ancestry on ‘*`Triglycerides`’* alongside their closest European neighbors. The bold study was used as representative study for the cluster, after semantic-based clustering. The colored column links to the color of the relative cluster in the UMAP displayed in (B). (B) Two-dimensional UMAP of the combined embeddings of European and African ancestry for the *‘`Triglycerides`’* cluster, showing similarities based on semantic similarity across GWAS of different ancestries. (C) Two-dimensional UMAP embeddings of GWAS traits based on semantic representations of the GWAS trait descriptions. The query trait (‘*`Alzheimer’s disease`*’) is represented in black. In addition, the five closest clusters based on cosine similarity are highlighted. (D) Semantic pairwise trait similarity heatmap based on cosine similarity, for the subset of traits shown in (C), illustrating cluster structure and redundancy among closely related phenotypes. (E) Genome-wide Miami-style overview combining haplotype-block level and SNP-level association signals for the representative GWAS of the Alzheimer’s disease cluster. This enables the comparison between the SNP association strength (uncorrected p-value <5×10^−8^) and the distribution of the number of haplotype-blocks containing at least one genome-wide significant variant (p<5×10^−8^), across chromosomes. All autosomal chromosomes 1-22 are shown. Sex chromosomes are omitted.

### Haplotype-level representation resolves the Alzheimer’s disease signal into 227 inherited units

To illustrate the trait-centric exploration, we queried *Alzheimer’s disease* (AD) within the *Haplotypes* module of snpXplorer. The platform retrieved 18 AD-related GWAS studies of European ancestry from OpenGWAS (*Table S3*). These included GWAS explicitly targeting AD as well as closely related phenotypes, such as maternal or paternal history of Alzheimer’s disease. Semantic similarity-based clustering grouped these traits together, and identified neighboring clusters associated with Parkinson’s disease (gold cluster), ADHD, autism spectrum disorder, and obsessive-compulsive disorder (green cluster), multiple sclerosis (cyan cluster), diseases of the nervous system and neuroticism (orange cluster), and macular degeneration (blue cluster) (*Figure 2C-D*). This similarity map thus highlights conceptual relationships among neurodegenerative and neuropsychiatric disorders.

The same *Haplotypes* module shows that for the selected representative AD GWAS [21], snpXplorer first identified 5,505 individual variants reaching genome-wide significance (p<5×10^−8^, *Figure 2E*, *Table S4*). These significant variants collectively map to 227 haplotype-blocks based on LD structure in European individuals (*Figure 2E* and *Table S5*). Interactive tables and plots enable the inspection of haplotype-block composition and the underlying variant associations, and all results can be downloaded for further investigation. Haplotype-block-level aggregation therefore groups the 5,505 significant AD variants into 227 inherited units, so that association evidence can be interpreted per haplotype-block rather than per individual variant. As an illustration, at the *APOE* locus, the largest common genetic risk factor for AD, the top variants rs429358 and rs7412, which jointly define the *APOE ε2/ε3/ε4* alleles, were assigned to two separate haplotype-blocks in both European and African haplotypes. Within a 2Mb region spanning rs7412 (43,908,822-45,908,822, GRCh38), snpXplorer identified 335 European and 869 African haplotype-blocks in total, reflecting the locus’s complex and extensive LD structure across ancestries as well as the deliberate choices of the LD threshold used. Of the 335 European haplotype-blocks, 69 contained 902/5505 SNPs significantly associated with AD (ebi-a-GCST90027158, p<5×10^−8^).

### A single TMEM106B haplotype-block carries synergistic pleiotropy across eleven neurological and behavioral traits

To showcase the gene-centric, cross-trait haplotype exploration, we focused on the *TMEM106B* locus, a gene associated with Alzheimer’s disease, frontotemporal dementia, and other aging-related phenotypes based on GWAS [1,22,23]. At the *TMEM106B* locus, snpXplorer identified twenty-two common haplotype-blocks in Europeans overlapping the gene region, with sizes ranging 35-345 kb (mean=182 kb, *Figure 3A-B-D, Table S6, and Figure S7*). Among haplotype-blocks containing at least one SNP with an association reported in OpenGWAS, distinct cross-trait association profiles were observed, ranging from broad to highly specific patterns of phenotypic association (*Figure 3C*). For example, haplotype-block B (7:12194394-12246599, *Figure 3A-B-C*) spans ∼52 kb, comprises 24 variants (LD structure shown in *Figure 3E*), and was associated with 8 distinct representative traits (see *Haplotype-block-level phenotype aggregation*), primarily related to neurological and behavioral phenotypes (depression, anxiety, tenseness, *Figure 3C*). This illustrates synergistic pleiotropic effects on nerve-related issues (ukb-b-6991) and depression (ebi-a-GCST006475), while antagonistic pleiotropy is reported on height (ukb-b-10787) (*Figure 3F*). Another haplotype-block (haplotype A, 7:12207704-12246424) of ∼38 kb in size contained 120 variants and was associated with 25 traits spanning neurological (Alzheimer’s disease, neuroticism, depression), metabolic (gastroesophageal disease, cholesterol), and cardiovascular conditions (hypertension, and diseases of the circulatory system). In contrast, snpXplorer identifies 74 common African haplotype blocks, with size ranging 1.4-459 kb (mean=115 kb, *Figure S7, Table S6*); only two of these reported previous associations from GWAS, consistent with the smaller sample sizes currently available for African-ancestry GWAS: haplotype-block A (7:12250352-12261908, 11.5 kb, 5 variants) is associated with leg predicted mass (ukb-e-23101), while haplotype-block B (7:12274144-12286905) is associated with childhood sunburn occasions (ukb-e-1737). A side-by-side visualisation of this region using European and African LD illustrates the finer-grained LD structure in African ancestry, with more numerous haplotype-blocks at *TMEM106B* locus (*Figure 3B, Table S6*).

**Figure 3:**
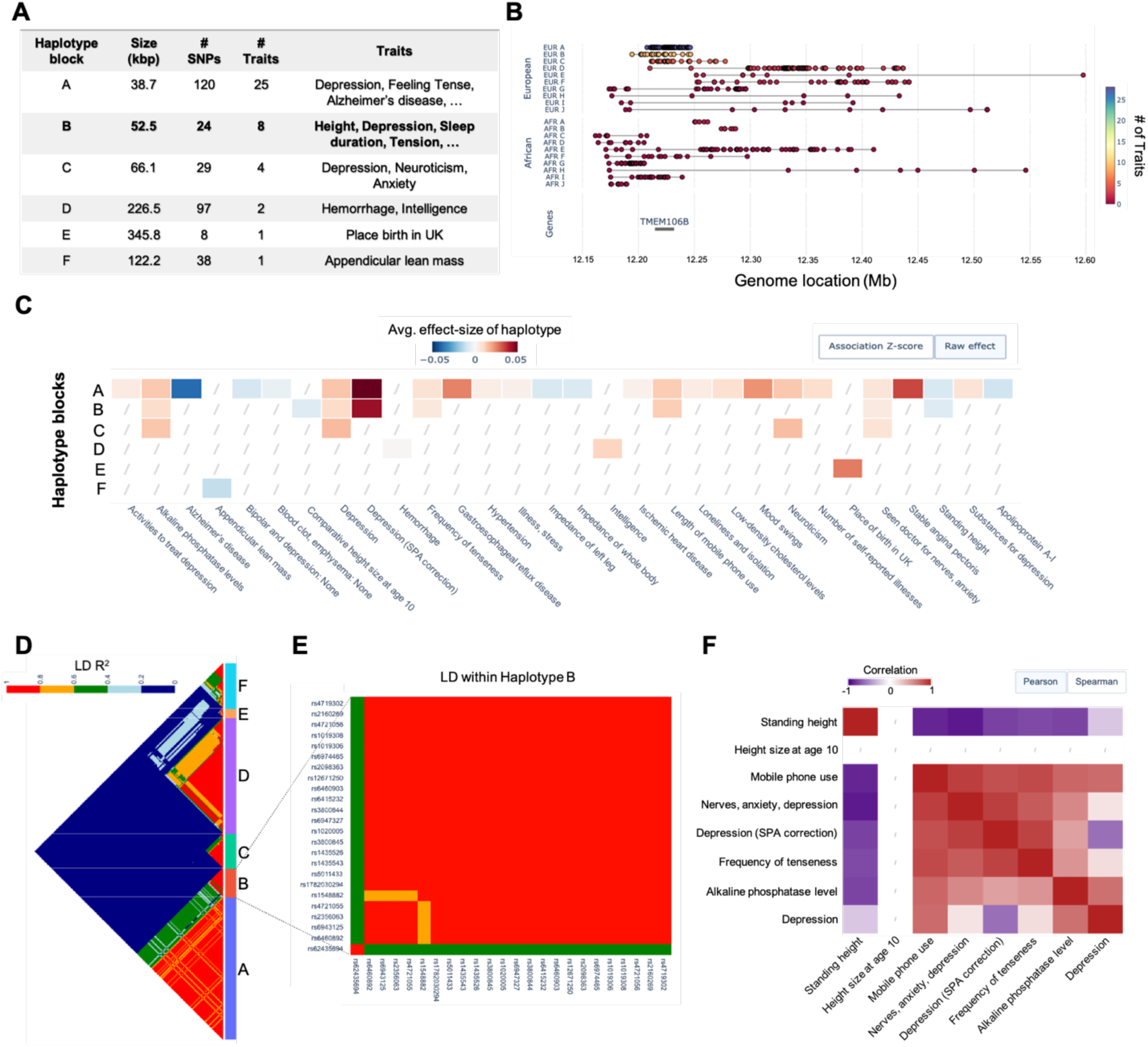
Gene-centric haplotype exploration for TMEM106B gene in snpXplorer. (A) Summary table reporting the haplotype-blocks identified in *TMEM106B* region for European ancestry with at least an association from OpenGWAS, including the relative number of SNPs, the number and name of the traits with associations of SNPs in the haplotype-blocks based on the OpenGWAS, and the size of the haplotype-block (kbp). Haplotype-blocks are sorted by the number of associated traits and given a letter as identifier. (B) Regional plot reporting the top haplotype-blocks in European and African ancestries sorted by the number of associated traits. Individual SNPs co-segregating within the same haplotype are connected on the same row, with the color representing the number of associated traits. (C) For each haplotype-block, the heatmap shows the average effect-size of SNPs within each block on phenotypes, based on SNP-associations from OpenGWAS. Only haplotype-blocks associated with at least one phenotype are shown. For visibility purposes, the top 40 phenotypes are shown. The user can choose whether the averaged effects reflect raw effect sizes from OpenGWAS or standardized (z-score) estimates. A maximum of 40 traits (columns) and 10 haplotype blocks (rows) are displayed. (D) LD patterns between all SNPs across haplotype-blocks in Europeans. The number of haplotypes can be dynamically adjusted from the user. LD R^2^ values are displayed. SNP Haplotype block membership is marked with the colored annotation column. (E) Focused LD heatmap showing all 24 SNPs and their pairwise LD scores in haplotype-block B (7:12194394-12246599). (F) Heatmap showing the trait-by-trait pairwise correlation of the effect of haplotype-block B. Positive and negative correlations reflect synergistic and antagonistic pleiotropic effects of haplotype-block B, respectively. Both Pearson’s or Spearman’s correlations can be shown (by default, Pearson’s).

### Haplotype-block context prioritizes PLEKHA1 at the chromosome 10q26 Alzheimer’s disease locus

To resolve the candidate gene at a second AD locus, we examined, as an illustrative example, rs7908662, an intronic variant near *PLEKHA1*. The variant, located in chromosome 10 at 122,413,396 (GRCh38, *Table S7*) has A as reference allele, and G, C, and T as alternative alleles, with G being the most common alternative allele. The G allele has an average frequency of 48% across all ancestries, ranging from 37% in African ancestry to 62% and 63% in East Asian and Finnish ancestries (*Figure 4A*).

**Figure 4:**
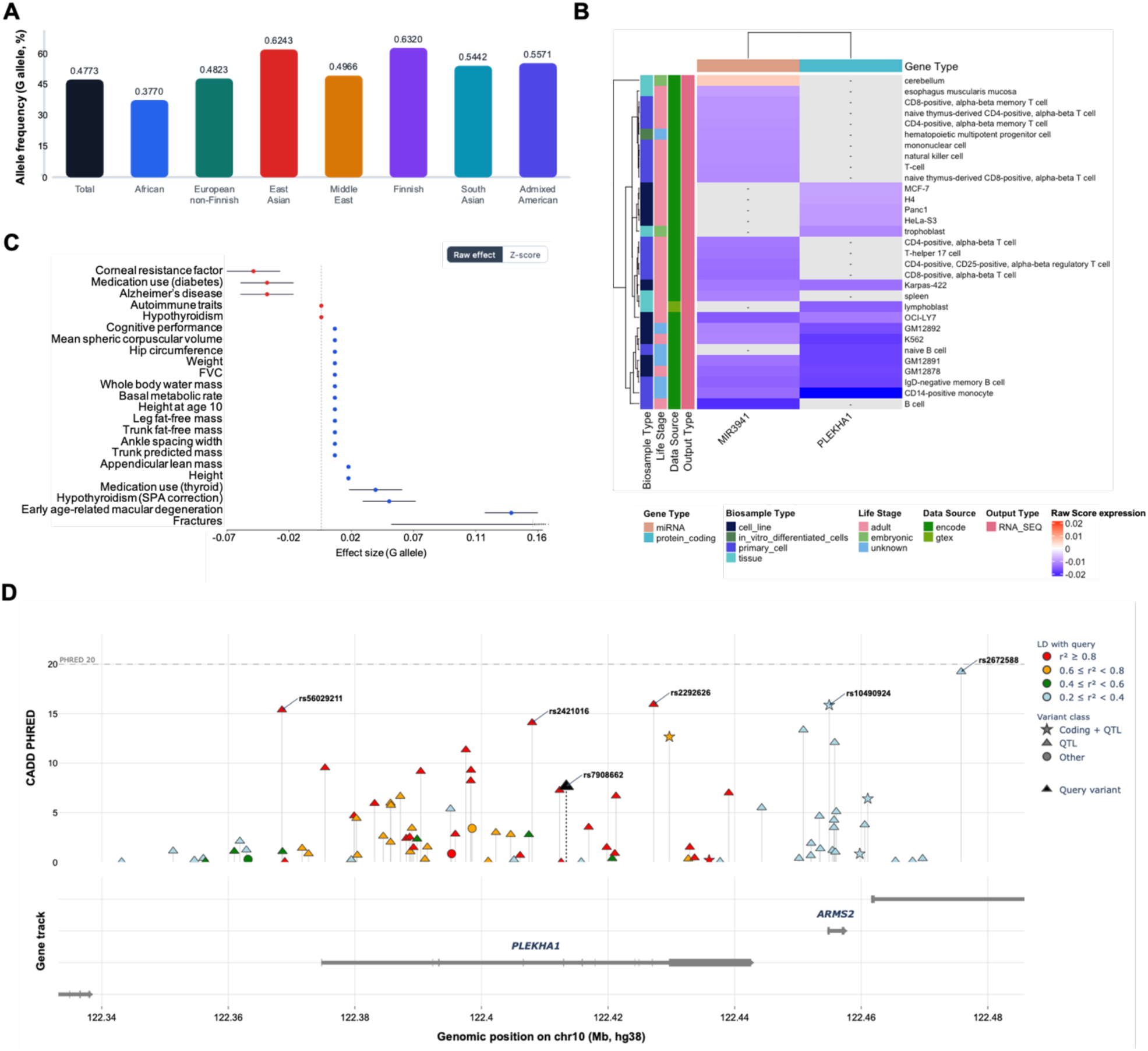
*Single variant annotation for rs7908662 within the PLEKHA1* gene. (A) Ancestry specific allele frequency plot showing the frequency of the alternative allele (G) across different ancestries, as reported in gnomAD. (B) Heatmap showing AlphaGenome annotation results. For visibility purposes, the top 0.5% annotations based on Quantile Score were selected. The raw score of these annotations is shown, with each row representing a specific single cell or a tissue. Side annotations report biosample type, life stages, data source, and output type (rows); and gene type (columns). The general pattern is that the alternative allele of the rs7908662 (G) results in a lower expression of both *PLEKHA1* gene and *MIR3941* miRNA. (C) Forest plot showing the effect of the alternative allele (G) of the variant on phenotypes from the OpenGWAS database. Red dots (effect size <0) refer to a decreased risk (for binary traits) or a lower value (for continuous traits) of the relative traits when carrying the G allele, while blue dots refer to an increased risk (for binary traits) or increased value (for continuous traits) for the relative traits. For example, the G allele is associated with lower risk of Alzheimer’s disease, and at the same time an increased level of cognitive performance. The user can choose whether to display raw effect sizes from OpenGWAS or standardized (z-score) effects. (D) Integrated visualisation of SNPs in the same haplotype-block as the target SNP alongside their features, including variant consequences, CADD score, and QTL annotation. Additional information including the specific tissues for QTL is displayed upon hovering over dots.

Functionally, rs7908662 is an intronic variant not annotated in ClinVar with a modest pathogenicity score (CADD=7.6), unlikely to be directly causal. QTL annotation indicated that the variant acts as an expression-QTL for *PLEKHA1* in multiple tissues, including brain (brain cerebellar hemisphere, effect_G_=−0.32, p=6.0×10^−8^), and also affects the expression of nearby genes in non-brain tissues (e.g. *ARMS2* gene in testis, effect_G_=−0.37, p=5.0×10^−16^) (*Figure 4D*). In line with this, AlphaGenome reports that the alternative allele of rs7908662 (G) is associated with changes in *PLEKHA1* and *MIR3941* expression across several immune- and brain-specific cell-types during both embryonic and adult life stage (*Figure 4B*). These also included brain-specific cell-types involved in Alzheimer’s disease mechanisms (*Table S8*). The variant also lies in a genomic region rich in structural variants content, including both short and long interspersed nuclear elements (SINE and LINE), with sizes up to 1.2 kb (*Table S9*).

Previous GWAS of European individuals have identified rs7908662 to be suggestively (p<5×10^−5^) associated with 25 traits (*Figure 4C*), including corneal resistance factor (ebi-a-GCST90100568, effect_G_=−0.05, p=3.6×10^−6^), autoimmune diseases (ebi-a-GCST90029015, effect_G_=−0.01, p=7.0×10^−6^), and Alzheimer’s disease (ebi-a-GCST90027158, effect_G_=−0.04, p=3.3×10^−6^). While the G allele has a protective effect on these traits, it shows antagonistic pleiotropy with other phenotypes including cognitive performance (ebi-a-GCST006572, effect_G_=0.01, p=2.5×10^−6^), age-related macular degeneration (ebi-a-GCST010723, effect_G_=0.14, p=5.7×10^−25^), weight (ebi-a-GCST90018949, effect_G_=0.01, p=4.1×10^−18^) and height (ukb-b-10787, effect_G_=0.02, p=5.7×10^−31^) (*Table S10*). No significant hits were found in GWAS of African ancestry, likely reflecting the smaller sample sizes currently available for African-ancestry GWAS at this locus. The above effect sizes are intended as raw estimates from OpenGWAS, however, standardized (z-score) effects can optionally be shown.

The European haplotype-block containing rs7908662 comprises 113 additional variants (LD R^2^>0.2, *Figure 4D*). Among these, snpXplorer identified both a non-synonymous variant (rs1045216, A>G, in *PLEKHA1,* LD R^2^=0.59) with a CADD score of 12.7; an intronic variant (rs2292626, C>T, LD R^2^=0.99) with a CADD score of 15.9 (*Figure 4D, Table S11*); and another non-synonymous variant with a CADD score of 15.9 in low LD (rs10490924, G>T, R^2^=0.21, *Figure 4D*). Both variant- and haplotype-block annotations were integrated to prioritize *PLEKHA1* as the most likely affected gene, while also highlighting secondary candidate genes supported by weaker LD or regulatory evidence (*ARMS2, MIR3941, HTRA1*). Assembled manually, the same evidence would require queries to at least five separate resources - CADD, gnomAD, ClinVar, GTEx and OpenGWAS - together with local LD computation against a reference panel, and several cross-referencing steps to align the resulting variant sets. snpXplorer returns this evidence within a single query. Integrative platforms reduce the number of primary sources a user must consult, but none combine these annotation layers with haplotype-block aggregation of association signals.

### PRS-ready variant sets

We have pre-calculated variant sets for all traits in the OpenGWAS, so that a PRS can be generated without uploading genotyping data, which is limited by privacy regulations. Using Alzheimer’s disease as an example, a manually curated set of variants as well as automated clumping-and-thresholding-based variant sets are available. It is possible to choose the variant set from the study of interest and at the p-value thresholds of interest (p<5×10^−5^, <5×10^−6^, <5×10^−7^, <5×10^−8^). Each downloadable variant set reports chromosome and position (GRCh38), effect allele and other allele, effect size and confidence intervals, p-value and allele frequency. These variant sets are fully compatible with external tools for PRS calculation (Jordan, PolyGenius), which can be downloaded and used to calculate PRS for the individuals of interest. snpXplorer also supports pathway-specific PRS: when the *Batch annotation* job is executed in combination with gene-set enrichment analysis, variant-pathway weights are automatically calculated. These weights estimate the effect of each individual variant on pathways [24], and can be directly used to calculate pathway-specific PRS with Jordan (*Figure S2C*).

### Responsiveness

To characterize responsiveness under typical use, we measured query latency for the single-variant annotation module and for the three haplotype query modes (haplotype blocks retrieved by variant, by gene, and by trait). For each module, 20 queries were selected at random and executed three times against the production server (https://snpxplorer.net), retaining the median latency per query. Median response times were 3.4s (IQR 1.8-6.6) for single-variant annotation, 1.0s (IQR 0.9-1.1s) for haplotype-block retrieval by variant, 1.1s (IQR 1.0-1.2s) by gene, and 10.6s (IQR 9.0-11.5) by trait. Retrieval by trait is slower than the other modes because it aggregates association signals across all haplotype-blocks of a study, rather than returning a single precomputed record.

## Discussion

snpXplorer is a freely available web platform that addresses important challenges in the interpretation of GWAS data by providing an integrated, haplotype-block-aware framework that combines large-scale GWAS resources with interactive visualization and multi-source annotation. This design allows users to click, browse, and inspect variants, haplotype-blocks, genes, and traits directly within the interface. As GWAS continue to expand in size, complexity, and genetic background, tools that enable efficient summarization, comparison, and contextualization of association signals are increasingly needed, and snpXplorer was developed to meet this demand. By enabling direct navigation between variants, haplotype-blocks, genes, and traits, snpXplorer streamlines GWAS and SNP interpretation workflows that would otherwise require multiple independent tools.

A central contribution of this new version of snpXplorer is its emphasis on haplotype-blocks, computed genome-wide for both European and African ancestry, laying the groundwork for cross-ancestry haplotype interpretation as African-ancestry GWAS resources continue to grow. While most existing variant interpretation tools remain focused on individual lead variants, haplotype-blocks can provide a more biologically informative representation of correlated genetic variation and facilitate the identification of affected genes that may be difficult to capture using SNP-centric approaches alone [1,2]. This integrated framework is particularly relevant for complex diseases, where GWAS loci often span multiple correlated variants and candidate genes. In these settings, interpretation based solely on individual lead SNPs may overlook broader regulatory mechanisms, which also include larger structural variants. Together, these features facilitate the interpretation of disease-associated loci within their broader genomic and functional context.

In addition, snpXplorer addresses the fragmentation of variant-level annotation across multiple resources. Existing platforms typically provide partial views of genetic evidence, requiring users to query multiple datasets independently. For example, ClinVar has a strong focus on clinically relevant variants, while most GWAS variants are not considered; gnomAD and dbSNP [25] are valuable for allele frequency and predicted consequences, but lack QTLs, haplotype information, and GWAS associations; phenome-wide association resources [26,27] report GWAS associations across traits but lack allele frequencies and QTL evidence; SNiPA [28] combines a rich variant annotation with GWAS associations, but remains primarily variant- and study-centric. In contrast, snpXplorer integrates clinical annotations, population allele frequencies, QTLs, LD structure, functional predictions, structural variants and GWAS associations within a unified interface. To the best of our knowledge, no single platform combines real-time multi-source variant annotation, haplotype-block context, and interactive cross-trait GWAS exploration within a single interface (*Table S12*).

snpXplorer facilitates cross-trait exploration by structuring large GWAS repositories using semantic similarity between phenotypes. This reduces redundancy and enables efficient navigation across related traits, supporting the identification of shared genetic architecture and pleiotropic effects. At the same time, the GWAS exploration module allows the visualization of multiple GWAS datasets simultaneously to compare association density tracks across traits.

Several widely used tools support GWAS and variant interpretation, including LocusZoom [29] for regional visualization, and platforms such as FUMA [30] and Open Targets Genetics [31] for GWAS downstream analyses, functional annotation and gene prioritization. Other resources, including PheWeb [32], PhenoScanner [33,34], and GWAS Atlas [35], facilitate querying of genotype-phenotype associations across studies. However, these tools typically rely on SNP-centric representations, provide static or list-based outputs, or cover only specific aspects of annotation. Rather than replacing these specialized resources, snpXplorer complements them by combining haplotype-block representation, integrated variant annotation, and interactive cross-trait exploration within a single framework.

Several limitations should be considered. snpXplorer relies on publicly available GWAS summary statistics and reference panels. The LD estimates used to define haplotype-blocks are currently based on individuals of European or African ancestry. Because LD structure, tagging patterns, and allele frequencies can differ substantially across ancestries, haplotype-block boundaries and the cross-trait associations reported here may not generalize to other populations. As with other GWAS interpretation tools, results should be interpreted in the context of the underlying datasets. Future developments will focus on incorporating additional ancestries (Asian and mixed ancestries), and will increase the supported annotation datasets. In addition, the clustering of traits based on semantic similarity heavily relies on OpenGWAS curation and the biomedical knowledge encoded in BioLORD and SapBERT rather than genetic correlation, [4,18,19] which needs genome-wide summary statistics and could be explored in future work. Nonetheless, semantic clustering surfaces relationships a keyword search would miss, such as grouping triglycerides with total cholesterol, LDL, and HDL levels, or prostate cancer with bladder and urinary tract cancers, pairs sharing no common vocabulary yet biologically related.

## Conclusion

In summary, snpXplorer provides an integrated platform for the exploration and annotation of GWAS and SNP data, combining haplotype-block-aware analyses with multi-source variant annotation, cross-trait integration, and support for both European and African ancestry. By addressing the fragmentation of existing tools and introducing haplotype-level representation, snpXplorer facilitates the interpretation of genetic association signals and supports a wide range of applications in human genetics research.

## Key Points

- snpXplorer represents GWAS association signals at the haplotype-block level, aggregating correlated variants into inherited units defined by recombination rate and pairwise linkage disequilibrium, rather than reporting individual lead variants.
- More than 11,000 GWAS datasets of European and African ancestry from OpenGWAS are integrated, enabling exploration across variants, haplotype-blocks, genes and traits without local upload of summary statistics.
- Variant interpretation is unified within a single interface, combining clinical annotations, cross-ancestry allele frequencies, functional predictions, quantitative trait loci, structural variation and reported GWAS associations, and extending each annotation to variants in linkage disequilibrium.
- Redundancy across closely related phenotypes is reduced by clustering trait descriptions with pretrained biomedical language embeddings, making navigation of large GWAS repositories tractable.
- Use cases in Alzheimer’s disease genetics show how haplotype-block-level representation exposes synergistic and antagonistic pleiotropy that variant-centric views can obscure.

## Acknowledgements

Not applicable.

## Author Contributions

Niccolò Tesi: Conceptualization, Formal analysis, Methodology, Validation, Writing of the original draft; Alex Salazar, Sven van der Lee, Gilad Green, Marc Hulsman, Henne Holstege: revision and editing of the original draft; Marcel Reinders: Conceptualization, revision and editing of the original draft.

## Ethics Approval

Genotyping data from 4,777 individuals of European ancestry were used for LD calculation to define haplotypes. All individuals provided written informed consent for the use of their genetic data for research purposes. The Medical Ethics Committee (METC) of the Amsterdam UMC approved all studies in which these individuals participated. Individuals of African ancestry (African American or Afro-Caribbean, Sub-Saharan African, and African unspecified) are publicly available from the 1000Genome project.

## Data Availability

All data used by snpXplorer are publicly available. snpXplorer is freely available at https://snpxplorer.net and https://snpxplorer.holstegelab.eu, and does not require user registration. The source code is available at https://github.com/TesiNicco/snpXplorer under an MIT license, with no restrictions on use by non-academics beyond those specified by that license. snpXplorer is implemented as a web application with a Python backend and an interactive frontend; analyses are performed server-side and results are rendered dynamically in the browser. The platform was tested in Safari, Google Chrome, Brave, Mozilla Firefox and Microsoft Edge. Genotyping data used for LD calculation are available upon reasonable request from the corresponding author. Documentation and code underlying the semantic similarity-based trait reduction and the haplotype block estimation are available at our https://github.com/TesiNicco/snpXplorer/tree/master/Exploration/accessory/haplotype_block_estimation_semantic_similarity.

## Supplementary

**Figure S1:**
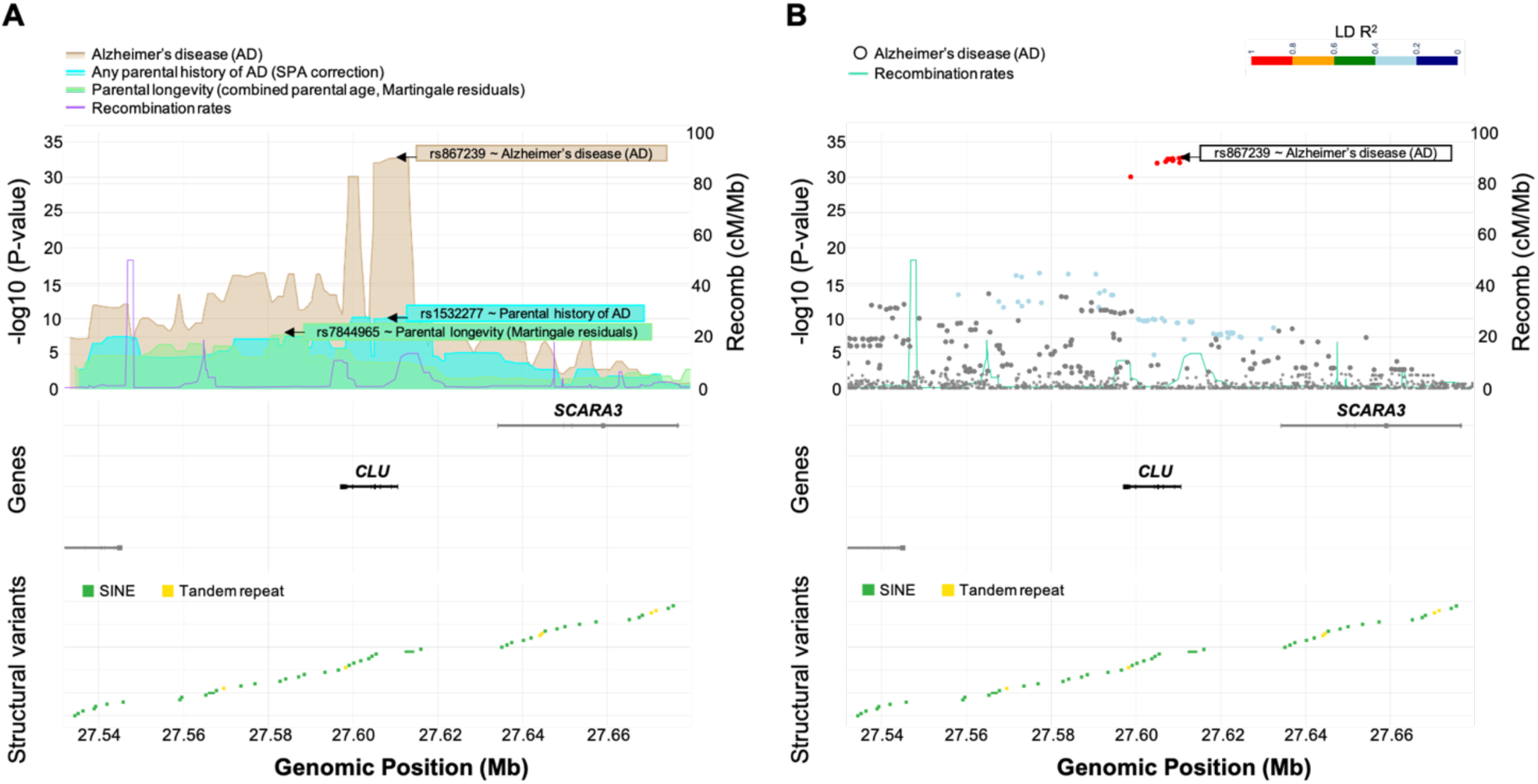
GWAS Exploration in snpXplorer: regional plots with gene and structural variant tracks. (A) Density track of significance level of multiple GWAS studies: GWAS of Alzheimer’s disease (ebi-a-GCST90027158, in brown); GWAS of parental history of Alzheimer’s disease (ebi-a-GCST90013971, in blue); GWAS of parental longevity (ebi-a-GCST006697, in green). The most significant variant for each study is annotated. Recombination rates are shown in purple as a single line. (B) A traditional locus plot of the same region in a single study (GWAS of Alzheimer’s disease), with dots indicating individual variants, and color indicating LD strength relative to the most significant variant in the region. Recombination rates are shown in grey as a single line. In both plots, the middle panel shows coding genes in the region, while the bottom panel shows structural variants in the region. As structural variants, tandem repeats (in yellow) and SINE elements (in red) are shown.

**Figure S2:**
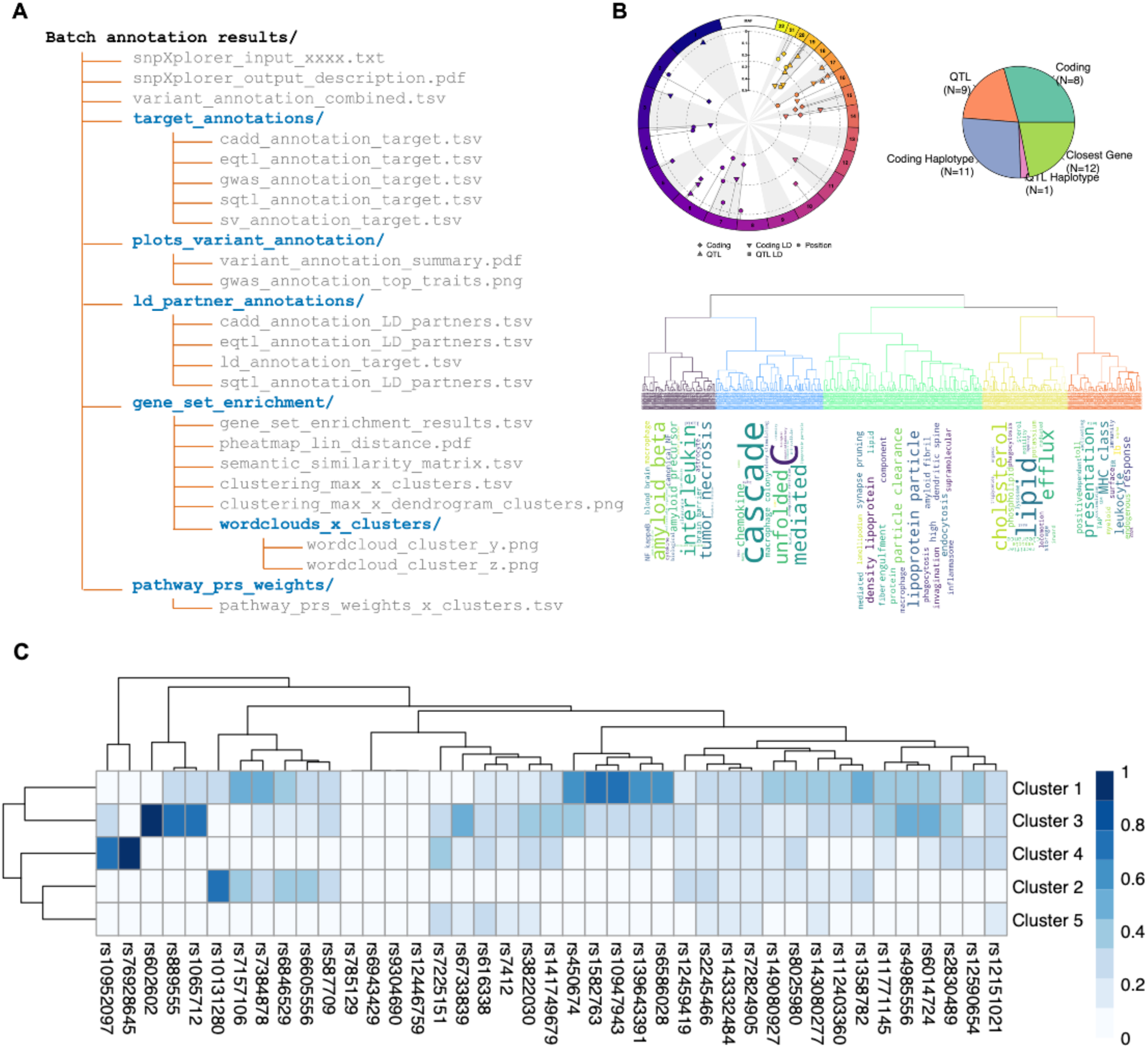
Batch annotation module. (A) The folder structure of the output data that is sent to the user when running the *Batch annotation* job. As input, up to 10,000 SNPs can be used for annotation and up to 1,000 SNPs can be used for gene-set enrichment analysis. A set of files including tables and graphs is saved and sent to the user, for further analyses. (B) Graphs showing the distribution of input SNPs across chromosomes, minor allele frequency, and annotation type (top left); a pie chart showing the proportion of coding SNPs, QTLs-SNPs, or SNPs annotated to the closest gene (top right). A dendrogram with relative word-clouds for each cluster showing the results of the gene-set enrichment analysis (bottom). Gene-set enrichment analysis is performed with gProfiler. Significant Gene-Ontology pathways are then clustered (agglomerative hierarchical clustering) based on semantic similarity. A dynamic cut-tree algorithm is applied to identify clusters of pathways. Word-clouds are finally used to provide functional context to each cluster. (C) Heatmap showing the variant-pathways mapping weights. These weights quantify the effect of SNPs on pathway-clusters, and allow the construction of pathway-specific Polygenic Risk Scores (PRS). A SNP is associated with a pathway depending on the associated genes, and the relative pathways. A score ranging 0-1 is assigned to each cluster, so that the cumulative effect of each variant across clusters is the same.

**Figure S3:**
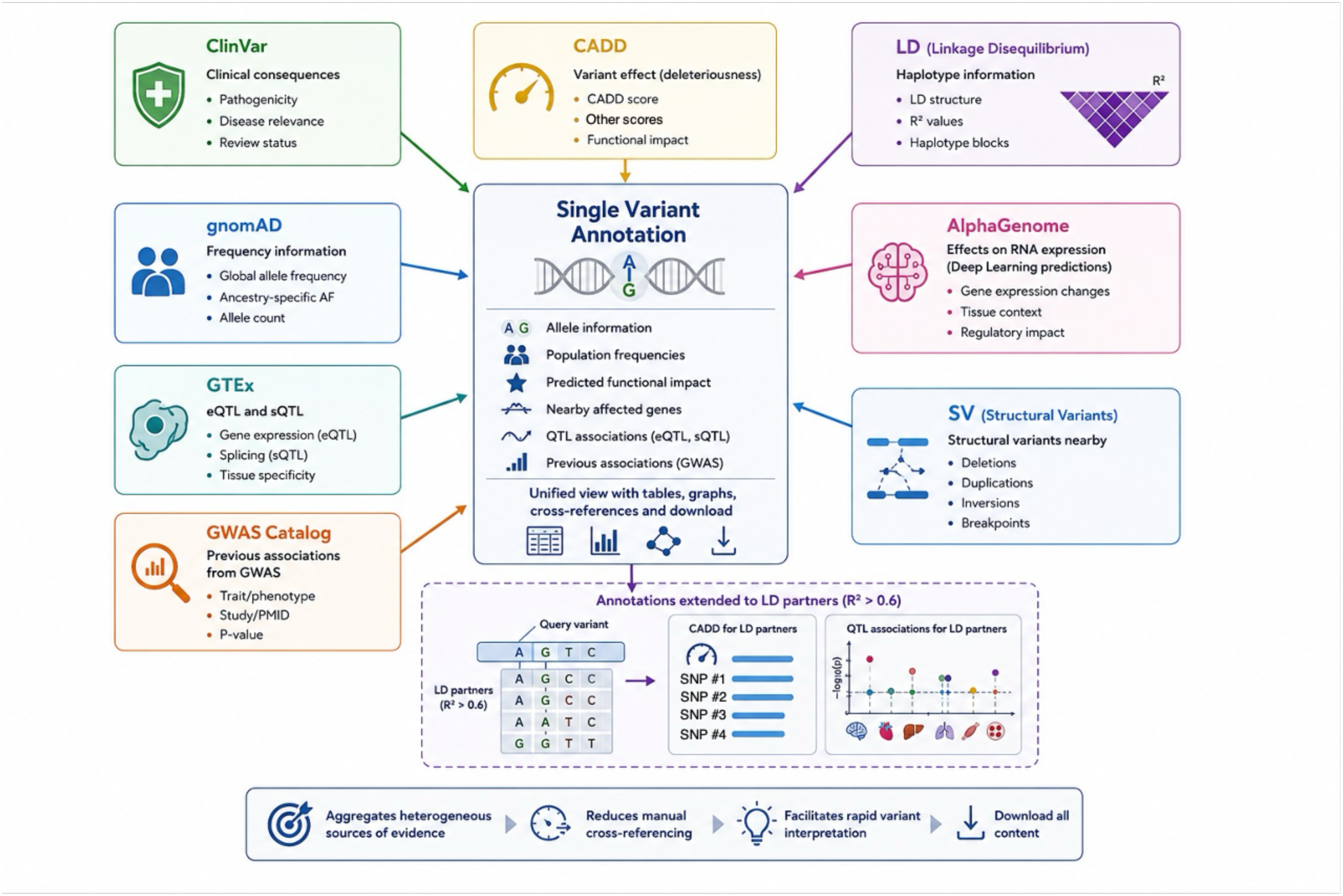
SNP annotation module. For a query variant, snpXplorer aggregates evidence from multiple data sources into a unified panel: clinical annotations from ClinVar, population allele frequencies from gnomAD, functional impact predictions from CADD and AlphaGenome, quantitative trait loci from GTEx, structural variant context, and previously reported associations from OpenGWAS. The module reports allele information, population frequencies, predicted functional impact, nearby affected genes, QTL associations, and GWAS associations, combined into tables, graphs, and cross-referenced links, with the option to download all content. Annotations are further extended to variants in linkage disequilibrium (LD, R^2^>0.6) with the query variant, allowing CADD scores and QTL associations of LD partners to be inspected alongside the query variant.

**Figure S4:**
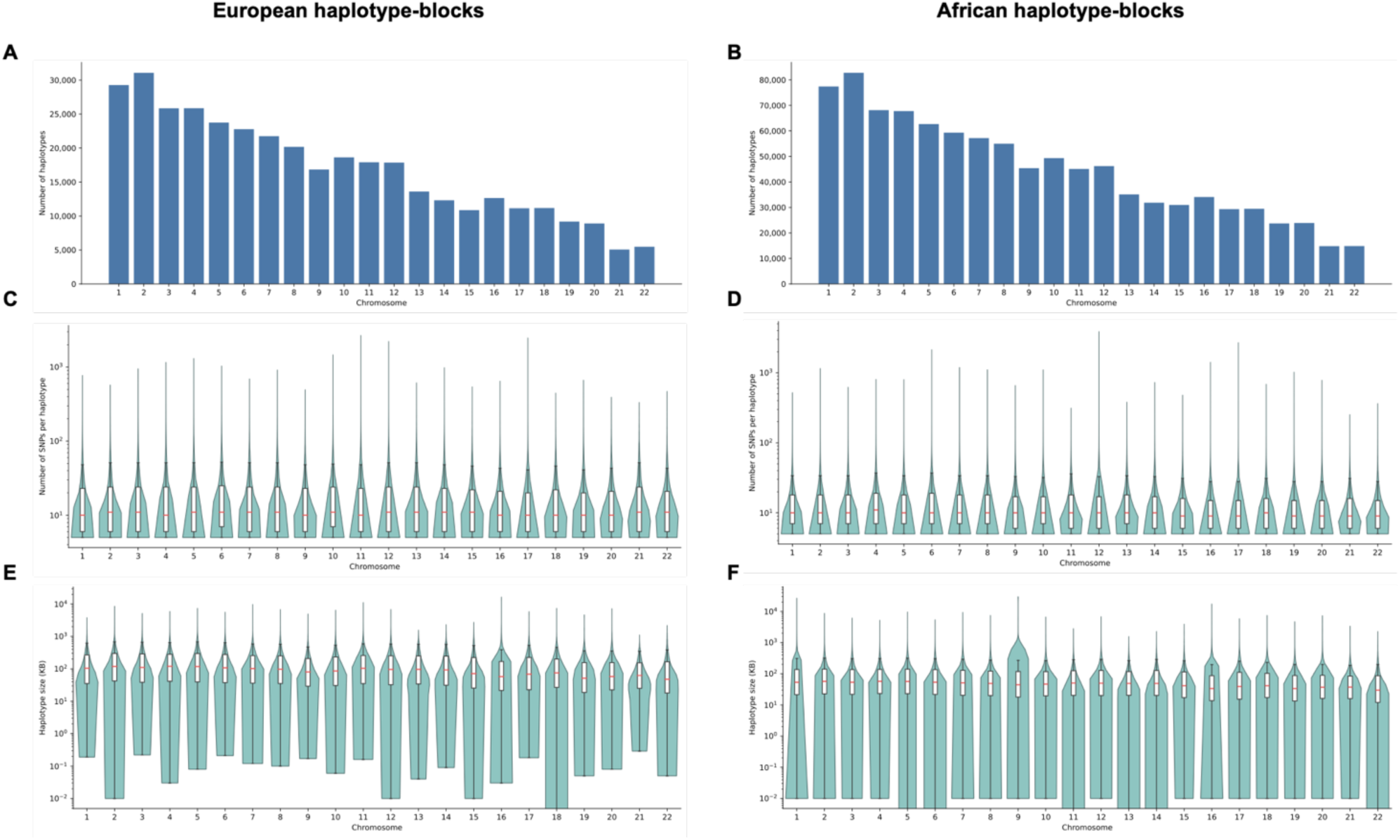
Haplotype block estimation in European and African individuals. Altogether, the haplotype blocks estimation revealed 372,097 common European haplotypes encompassing 8,225,502 SNPs, and 984,474 common African haplotype blocks comprising 15,404,984. The higher number of African haplotype blocks and variants can be due to the higher heterogeneity of African samples, and the nature of sequencing data compared to imputed genotyping data. (A, B) The number of common haplotypes across chromosomes, respectively in European (A) and African (B). (C, D) The distribution of the number of SNPs per haplotype, across chromosomes, respectively in European (C) and African (D). (E, F) The distribution of the haplotype size (in kilo base pairs, KB), across chromosomes, respectively in European (E) and African (F).

**Figure S5:**
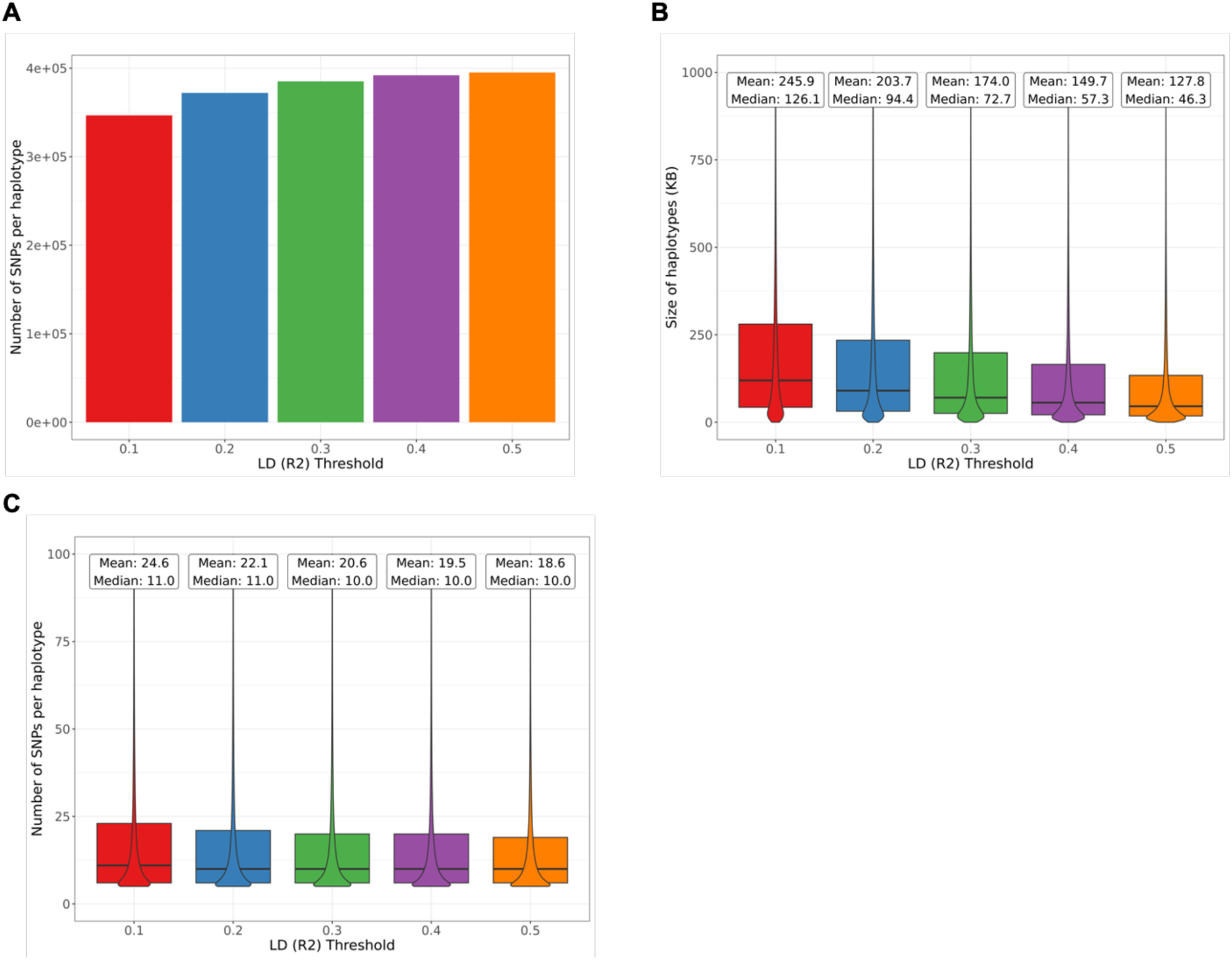
Sensitivity analysis of LD thresholds chosen for defining haplotype blocks. (A) Shows the number of haplotype-blocks across LD R^2^ thresholds. (B) Shows the size of haplotype-block alongside mean and median values across LD R^2^ thresholds. (C) Shows the number of SNPs per haplotype-block alongside mean and median values across LD R^2^ thresholds. For the final set of haplotype-blocks we used a threshold of LD R^2^=0.2. This analysis was performed in European individuals only.

**Figure S6:**
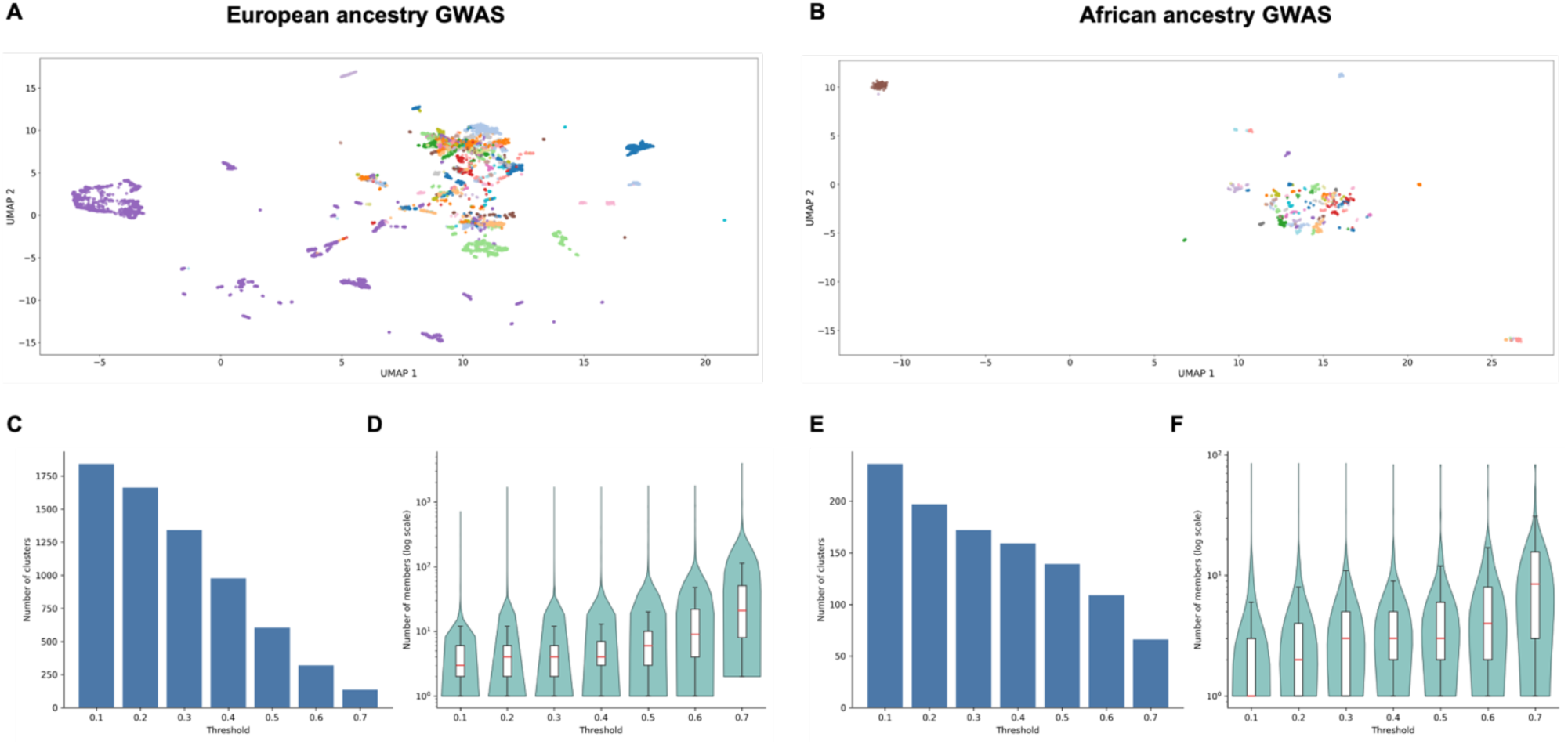
Semantic-based clustering of GWAS traits across European and African ancestries. Altogether, semantic clustering of 10,604 European and 751 African GWAS trait descriptions yielded 604 and 136 clusters (respectively for European and African) at the selected cosine distance threshold of 0.5. (A, B) UMAP embedding of the combined BioLORD-2023 and SapBERT trait embeddings, with points colored by the cluster assigned at a cosine distance threshold of 0.5, respectively for European and African traits. (C, E) The number of clusters obtained as a function of the cosine distance threshold, ranging from 0.1 to 0.7, respectively for European and African traits. (D, F) The distribution of the number of traits per cluster, across thresholds, respectively for European and African traits.

**Figure S7:**
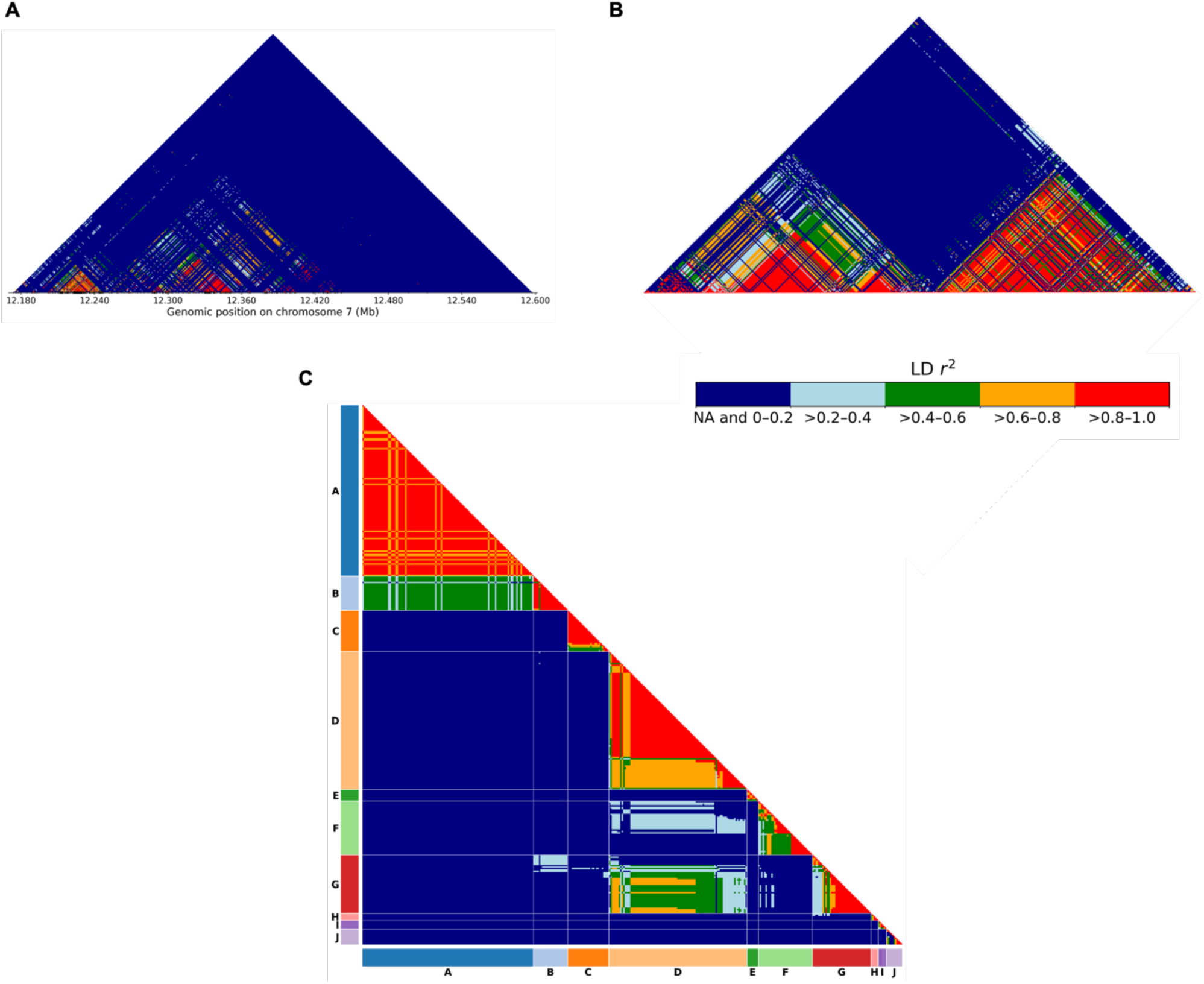
Combined representation of haplotype-blocks at TMEM106B gene. For all figures, the top 10 haplotype-blocks based on their number of associations are shown. (A) Shows the haplotype-block LD patterns (R^2^) with constrained physical position. (B) Shows the haplotype-block LD patterns restricting to SNPs in the region and ordered by position. (C) Shows the haplotype-block LD patterns restricting to SNPs in the region and ordered by haplotype-block, and within each block, by position

